# Genetics of growth rate in induced pluripotent stem cells

**DOI:** 10.1101/2025.07.02.662844

**Authors:** Brian N. Lee, Henry J. Taylor, Filippo Cipriani, Narisu Narisu, Catherine C. Robertson, Amy J. Swift, Neelam Sinha, Tingfen Yan, Lori L. Bonnycastle, Nathan Dale, Annie Butt, Hemant Parsaud, Stefan Semrau, NYSCF Global Stem Cell Array Team, GENESiPS Consortium, iPSCORE Consortium, Joshua W. Knowles, Ivan Carcamo-Orive, Agnieszka D’Antonio-Chronowska, Kelly A. Frazer, Leslie G. Biesecker, Scott Noggle, Michael R. Erdos, Daniel Paull, Francis S. Collins, D. Leland Taylor

## Abstract

Human induced pluripotent stem cells (iPSCs) have transformed biomedical research by enabling the generation of diverse cell types from accessible somatic tissues. However, certain fundamental biological properties, such as the genetic and epigenetic determinants of iPSC proliferation, remain poorly characterized. We measured the growth of iPSC lines derived from 602 unique donors using high-throughput time-lapse imaging, quantified proliferation through a growth Area-Under-the-Curve (gAUC) phenotype, and correlated gAUC with the gene expression and genotype of the cell lines. We identified 3,091 genes associated with gAUC, many of which are well established regulators of cell proliferation. We also found that rare deleterious variants in *WDR54* were associated with reduced iPSC growth and that *WDR54* was differentially expressed with respect to gAUC. Although no common variants showed a genome-wide association with gAUC, iPSC lines from monozygotic twins were highly correlated, and common genetic variation explained approximately 71-75% of the variance in iPSC growth rates. These results indicate a complex genetic architecture of iPSC growth rates, where rare, large-effect variants in important growth regulators, including *WDR54*, are layered onto a highly polygenic background. These findings have important implications for the design of pooled iPSC-based studies and disease models, which may be confounded by intrinsic growth differences.

## Introduction

The development of human induced pluripotent stem cells (iPSCs) has transformed many aspects of biomedical research. The first iPSCs were described 18 years ago, derived from human fibroblasts reprogrammed into a pluripotent state.^1–3^ Since then, numerous protocols have been established to differentiate iPSCs into a wide variety of cell types, including those difficult to obtain as primary human tissues.^4–8^ Given this differentiation potential and the facility of *in vitro* culture and manipulation, iPSCs offer a promising avenue for a wide array of basic science investigations, as well as translational programs including testing of drug and cellular therapies^9,10^ and potential autotransplantation.^11^

Many basic biological questions that are difficult to address *in vivo* can be assessed with iPSCs. For example, disease-associated loci discovered through genome-wide association studies (GWASs) often fall in non-coding regions of the genome, masking the underlying effector gene(s) and the specific cellular mechanisms underlying disease risk.^12,13^ Studies examining the genetics of gene expression across various tissues have not resolved all of these associations, suggesting that some disease signals may originate from cell types or cell states that are difficult to capture *in vivo* (e.g. pancreatic progenitor cells or beta cells after glucose stimulation).^14^ Large-scale genetic studies using iPSCs offer one way to investigate the effects of natural human variation and disease-associated variants in otherwise inaccessible cell types and contexts.^15–19^ As repositories of iPSCs derived from many donors continue to expand, such studies will likely become commonplace, making a detailed understanding of iPSC properties necessary to optimize study designs and account for confounding factors.

In this study (Figure 1), we considered one such variable—growth rates of undifferentiated iPSCs. Growth rates can be a critical factor in pooled study designs where different proliferation rates may cause certain lines to dominate a pool.^15,20^ Across 602 lines, we measured rates of cell growth from daily images, paired with RNA sequencing on a subset of 333 lines. Using these data, we performed an association analysis of iPSC growth rate with gene expression, common genetic variation, and rare genetic variation. Our results support a polygenic model of cell proliferation, whereby common variants contribute small effects and rare variants may exert larger influences on cellular growth dynamics.

**Fig. 1.**
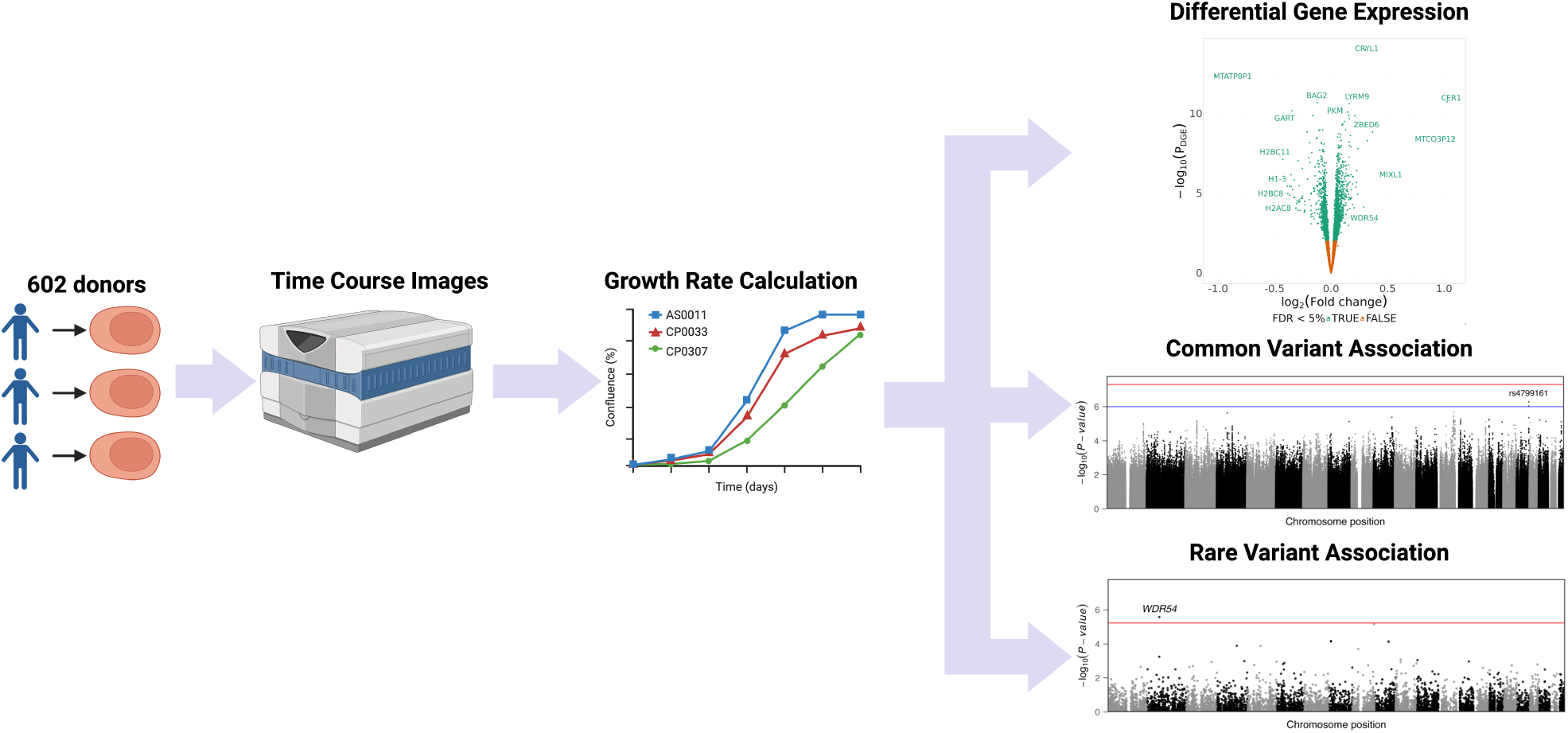
Graphical overview of this study. Human iPSCs from multiple cohorts were cultured in 96-well plates until confluence. Daily cell density measurements were obtained via imaging cytometry. Upon reaching confluence, cells were harvested for bulk RNA sequencing, genotyping, and genome sequencing, enabling differential expression analysis and both common- and rare-variant association studies.

## Results

### Cohort characterization and overview of genetic data

We obtained iPSC lines from 602 donors (ESM Table 1).^21–26^ These lines included both male (278 lines; 46.2%) and female (324 lines; 53.8%) donors. Lines were grown in two types of media, either mTeSR Plus (n = 347) or Freedom (n = 255), which we controlled for in subsequent analyses as a covariate (Methods).

We genotyped all 602 donors and imputed variants to GRCh38 (Methods). Focusing on common single nucleotide polymorphisms (SNPs; minor allele frequency [MAF] > 5%), we retained 5,985,990 variants after quality control procedures. To explore the genetic diversity represented in the full cohort, we grouped participants according to their genetic similarity to 1000 Genomes genetic ancestry groups, including participants from European (EUR), African (AFR), American (AMR), East Asian (EAS), and South Asian (SAS) ancestry groups (Methods).^27^ For downstream analyses, we described the genetic similarity groups as 1KG-EUR-like, 1KG-AFR-like, 1KG-AMR-like, 1KG-EAS-like, and 1KG-SAS-like, respectively. Most of our participants were 1KG-EUR-like (n = 446), but we also identified participants that were 1KG-AFR-like (n = 39), 1KG-AMR-like (n = 44), 1KG-EAS-like (n = 51), and 1KG-SAS-like (n = 22) ancestry groups (S. Fig 1).

While genotype imputation is accurate for common SNPs, rare genetic variants are often missed. To identify rare genetic variants, we used genome sequencing (GS) or exome sequencing (ES) data to call exonic variants in 514 donors (Methods). For the majority of the cell lines, DNA originated from the iPSC lines; for a subset (n = 195), however, we used DNA obtained directly from donor cells, either blood or fibroblasts. On average, we generated approximately 656.0 million reads per cell line for a depth of 31X. After quality control procedures, we retained 178,745 exonic single nucleotide or short insertion-deletions variants for downstream analyses.

### Derivation of growth rate from cell images

To measure cell growth rate, we plated cells from the iPSC lines of all 602 donors (ESM Table 1) on the New York Stem Cell Foundation (NYSCF) robotic culture system, performed at least one passage, and seeded cells at a standard density on fresh plates. We used a Celigo imaging instrument to image each well once a day from which we measured well coverage (expressed as a percent). In total, we had 139,246 timestamped well coverage measurements across 26,066 wells spanning all 602 lines (average 43.3 wells per cell line). For each well, we used the series of well coverage measurements to derive a growth Area-Under-the-Curve (gAUC) to represent growth rate for downstream analyses (Methods). After quality control procedures (Methods), we retained on average 27.0 wells with gAUC estimates per line (25.7 standard deviation [SD]) and observed high reproducibility across the replicate wells (intraclass correlation coefficient [ICC] = 0.96, *P* = 1.11×10^−16^, F-test). We also compared gAUC in 16 twin participants (eight pairs) and found them to be correlated (ICC = 0.65; *P* = 0.02, F-test). These observations suggest a genetic component of gAUC, though cell-specific effects or technical artifacts may also contribute (e.g., epigenomic events that take place during the development of pluripotency from a differentiated cell). To explore the effect of known technical or biological factors on gAUC, we tested for gAUC association with a variety of iPSC attributes including genetically-imputed donor sex, pluripotency reprogramming method, starting cell type for reprogramming, and cell culture media used (Methods). We found three terms to be associated with gAUC: cell culture media (*P* = 0.008), starting cell type (*P* = 1.68×10^−15^), and pluripotency reprogramming method (*P* = 1.80×10^−7^) (Fig 2A).

**Fig. 2.**
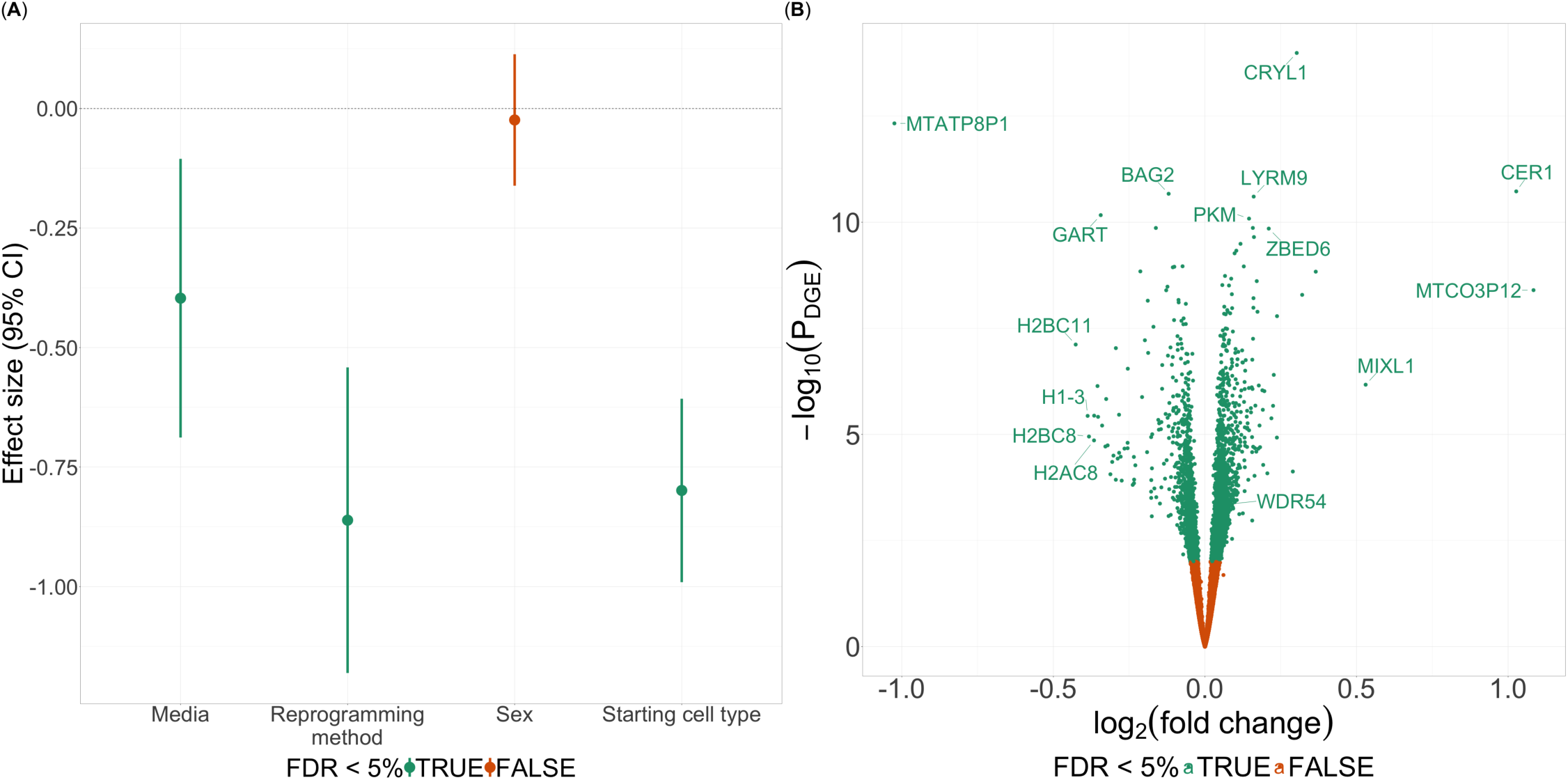
Covariate associations with growth Area-Under-the-Curve (gAUC) and differential gene expression analyses. (A) Effect size (y-axis) with 95% confidence intervals (CIs; lines) of iPSC line characteristics (x-axis). Color denotes FDR *<* 5%. (B) Volcano plots of *log*_2_(fold change) (x-axis) and *→log*_10_(*P*) (y-axis) from differential gene expression analysis. Color denotes associated genes (FDR *<* 5%).

### Growth-associated differential gene expression analysis

We sought to characterize the transcriptional profile of iPSC growth rates and generated RNA-sequencing data for 333 lines after the lines were standardized to a uniform mTeSR Plus media (Methods). We generated 235 million reads on average. After quality control and gene expression quantification, we retained expression data for 15,603 genes (Methods).

We tested for associations of gene expression with gAUC and identified 3,091 differentially expressed genes (DEGs; false discovery rate [FDR] < 5%; Fig. 2B; Methods). Many of the genes that we found are well established cell growth and proliferation genes, confirming the gAUC phenotype: *KDM4C*,^28^ *FGF8*,^29^ *SLITRK1*,^30^ *NRCAM*,^31^ *L1CAM*,^32^ *COBL*,^33^ *SYT4*,^34^ *KRT17*,^35^ *NCAPG2*,^36^ *PIK3CA*,^37^ *AXIN2*,^38^ and *RNF43.*^39^

### Contribution of common genetic variants to growth rate

We performed a genome-wide common variant association study (CVAS) to test for common variant associations with gAUC using all 602 iPSC lines, controlling for culture media type, starting cell type, reprogramming method, and the first ten genetic principal components (PCs) as covariates (Methods). We did not identify any genome-wide associated signals (*P* < 5×10^−8^). However, we found one locus with suggestive evidence of association (*P* < 1×10^−6^; Fig 3A), tagged by rs4799161. The rs4799161 variant is located within the *RP11-383C5.4* lncRNA and is approximately 451 kb upstream of *SALL3*, the nearest protein-coding gene. *SALL3* has been found to influence methylation of Wnt signaling-related genes in iPSCs and, therefore, may play an important role in both cell proliferation and stem cell differentiation pathways.^40^

**Fig. 3.**
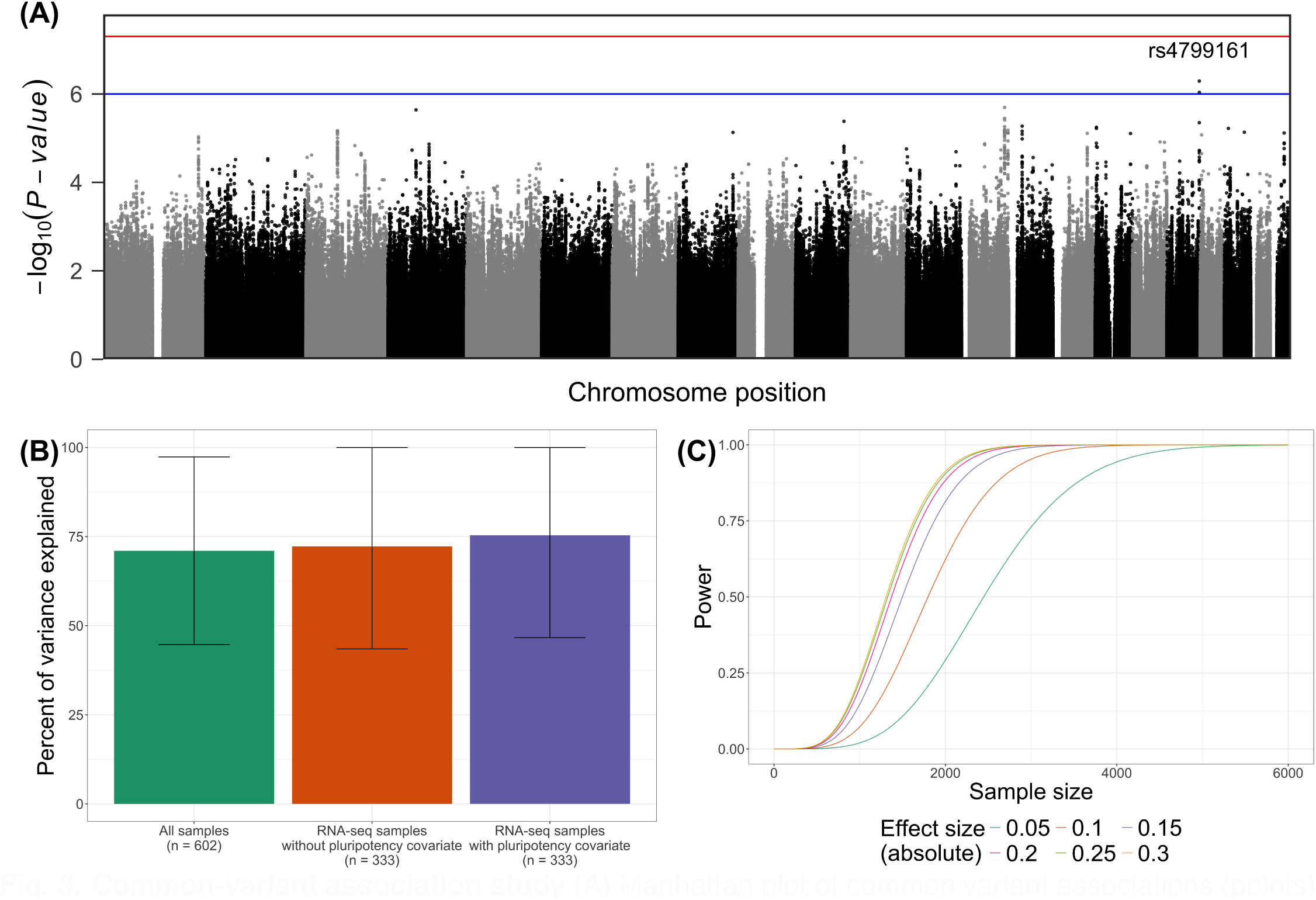
Common-variant association study. (A) Manhattan plot of common variant associations (points) with gAUC. Nominal association threshold (*P <* 1 *↑* 10*^→^*^6^) is indicated by the blue line. Genome-wide association threshold (*P <* 5 *↑* 10*^→^*^8^) is indicated by the red line. (B) Percent of growth rate variance explained by the genetic relatedness matrix (y-axis) across different models (x-axis). The horizontal axis and colors indicate the model—either all samples (green), samples with RNA-seq (orange), or RNA-seq samples with pluripotency covariates (purple). The vertical error bars indicate 95% confidence intervals (CIs). (C) Power estimates (y-axis) by sample size (x-axis). Color indicates absolute effect size modelled.

Though we identified limited common variant associations, a substantial genetic component to gAUC may exist, consisting of highly polygenic, small effect sizes. To better understand the genetics of gAUC, we estimated the SNP-based, narrow-sense heritability modelling the genetic relatedness matrix (h_g_ ^2^) along with pluripotency of lines when applicable (Methods). We analyzed the heritability in three ways: (i) across all samples, including media, starting cell type, and reprogramming method as covariates (n_All_ = 602), (ii) across lines with RNA-seq, where we used expression of *POU5F1*, *PODXL*, *NANOG* and *SOX2* to control for the pluripotency (n_RNA_ = 333; only starting cell type and reprogramming method as covariates since all of the RNA-seq lines were grown in mTeSR media), and (iii) across lines with RNA-seq without controlling for pluripotency. Across all models, we found that the genetic component explained approximately 71-75% of the variance in gAUC, indicating a strong heritable component to iPSC growth rates (Fig. 3B). These findings are consistent with the high intraclass correlation coefficient identified within twin cell line pairs.

Finally, given our heritability findings, we hypothesized that the limited CVAS results were a consequence of an insufficient sample size. We performed calculations to determine an optimal sample size for investigating the effects of common variants on iPSC growth rates (Methods). We estimated that identifying common variants associated with growth phenotypes at 80% power would require a minimum sample size of 1,720 individuals to detect effect sizes of 0.30 and up to 3,222 individuals to detect smaller effects of 0.05 (Fig. 3C).

### Contribution of rare genetic variants in protein coding regions to growth rate

For a highly polygenic phenotype, a CVAS often identifies loci with small effect sizes.^41^ However, rare deleterious or *de novo* mutations in coding regions may have large effects on cellular growth, as is evident in many cancers.^42^ To explore the role of moderately rare and rare variants (minor allele frequency [MAF] < 1%) in gAUC, we performed a rare variant association study (RVAS) across 514 donors with GS or ES by aggregating protein-coding variants across genes and testing gene-based burden scores with gAUC (Methods). We considered only variants with a predicted functional consequence of loss of function (pLoF) or putatively damaging missense with a CADD score ≥ 20. In total, after additional quality control filters (Methods), we tested for associations of 8,480 genes with gAUC.

We identified one association with decreased gAUC at *WDR54* (*P*_Bonferroni_ < 0.05; Fig. 4A), driven by five potentially damaging missense variants in *WDR54* (ESM Table 2). We did not detect any pLoF variants in *WDR54*. Because these missense variants occurred in DNA derived from both donor fibroblasts (n = 6) and iPSC lines themselves (n = 11), we cannot determine if they are somatic or germline. Given the difficulty in predicting the pathogenicity of missense variants, we conducted a sensitivity analysis by repeating the RVAS using a stricter CADD score threshold (≥ 25), resulting in 3,441 tests. Although the number of missense variants considered in *WDR54* decreased to two, the association with decreased gAUC remained (*P*_Bonferroni_ < 0.05), supporting the robustness of this finding. We did not identify any additional associations in this restricted analysis.

**Fig. 4.**
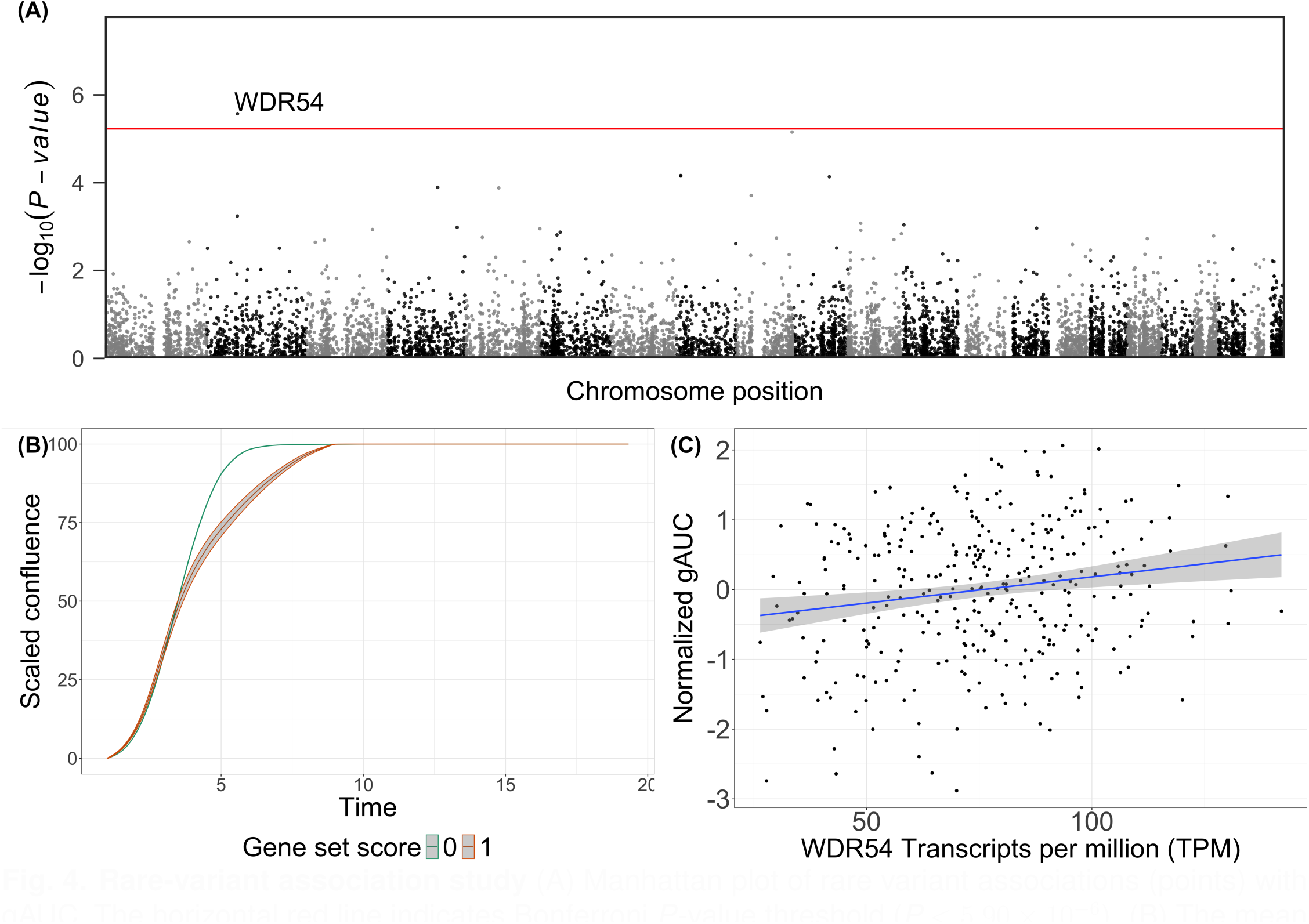
Rare-variant association study. (A) Manhattan plot of rare variant associations (points) with gAUC. The horizontal red line indicates Bonferroni *P*-value threshold (*P <* 5.90 *↑* 10*^→^*^6^). (B) The mean confluence trajectories stratified by rare-variant burden scores (color) for *WDR54* with confluence percentage (y-axis) over time (x-axis). (C) A scatterplot displaying the relationship between *WDR54* gene expression (x-axis) and the average gAUC (y-axis) per line (points). The shaded ribbon indicates the 95% CI.

Finally, to further explore the *WDR54* rare variant association, we reconsidered the gAUC differential gene expression results and found an association of *WDR54* expression with gAUC (FDR < 5%; *P* = 4.58×10^−4^, Fig. 2B, 4C). Combined with the rare variant findings, these findings are consistent with previous reports of increased expression of *WDR54* and cell/tumor proliferation.^43–45^

## Discussion

In this study, we investigated iPSC growth rates across 602 unique donors using a unique robotic culture system and an imaging instrument that allowed highly reproducible daily capture of cell growth. From the RNA-seq analysis, we identified thousands of differentially expressed genes associated with growth rate. Our rare variant association analysis identified *WDR54* as having putatively damaging missense variants associated with reduced growth rate. This finding was also corroborated by our RNA-seq results, where increased expression of *WDR54* was associated with increased growth rate. Our common variant association analysis did not identify variants associated with iPSC growth (*P* < 5×10^−8^), which our power analysis shows will require larger sample sizes.

Since we studied iPSC lines derived by different laboratories using different methods, it was reasonable to consider that other epigenetic factors might influence the growth rates of individual lines. We pursued three approaches to address this question: (i) We considered 16 cell lines from eight pairs of monozygotic twins and found their growth rates to be highly correlated (ICC = 0.65), though not as high as replicates of the same cell line studied at different times (ICC = 0.96). (ii) We decomposed the variance explained by a genetic relatedness matrix and observed high heritability, approximately 71-75%. (iii) We searched for and found differences in growth rates of iPSCs depending on the starting cell type (blood or skin), the pluripotency reprogramming method (mRNA-based or Sendai virus), and the media used for cell culture (mTeSR Plus or Freedom). Our study accounts for these differences by adding these three variables as covariates in our statistical analyses. These findings demonstrate that, after accounting for methodological sources of variation, a substantial proportion of the remaining variance in iPSC growth is attributable to genetic factors.

The differential gene expression analysis identified numerous genes with expression differences associated with growth rates. Many gAUC-associated genes are known oncogenes including *FGF8*,^29^ *L1CAM*,^32^ *KRT17*,^35^ *NCAPG2*,^36^ *PIK3CA*,^37^ and *RNF43.*^39^ The differential expression of genes associated with chromatin remodeling (i.e. *KDM4C*),^28^ cytoskeletal reorganization (*COBL*),^33^ and DNA damage repair (*KDM4C*)^28^ further implicates cellular features that modulate cell growth.

Finding the *WDR54* association with iPSC growth rates comports with prior studies of the function of this gene. *WDR54* is a prominent oncogene within colorectal cancer, bladder cancer, and T-cell acute lymphoblastic leukemia, and has also been proposed as a therapeutic target for head and neck squamous cell carcinoma.^44–46^ Knockdown of *WDR54* causes reduced cell growth and increased apoptosis.^43,44,46^ Mechanistic studies show that *WDR54* enhances extracellular signal-regulated kinase activation—a process that ultimately regulates cell growth, proliferation, and differentiation—by sustaining epidermal growth factor receptor signaling at the plasma membrane.^47^ These established roles of *WDR54* in cellular proliferation reinforce the biological context for the gene expression and rare-variant association finding.

Strengths of the study include the large number of lines considered, an automated system to precisely measure cell growth over time, and a broad characterization of iPSC growth contributors through SNP genotyping, DNA sequencing by ES and GS, and bulk RNA sequencing. One limitation is the inability to distinguish germline from somatic mutations in these iPSC lines; future studies may benefit by identifying somatic mutations through comparison of iPSC sequence with blood or skin DNA sequence from the donors.^48^ A second limitation is the inability to detect common variants associated with growth rate, suggesting future studies may need to be several times larger to have the necessary power to detect these polygenic effects. A third limitation is our dependence on computational methods to identify pathogenic variants in the coding region of genes, which may improve in the future with a combination of experimental work and more advanced artificial intelligence models of protein structure and variant function.

Awareness of the genetic drivers of iPSC growth can provide important insights for future research designs. For instance, in cell village association studies, lines with aggressive growth and enhanced differentiation characteristics can overwhelm a heterogeneous pool of cells, potentially confounding experimental outcomes.^15,16,20^ As large-scale genetic studies of iPSCs become more common, study designs should consider differences in starting cell type, type of media, and pluripotency reprogramming method. These studies may also benefit from genetic screening of individual lines for both somatic and germline rare variants in relevant genes, including *WDR54*.

## Extended authorship

**NYSCF Global Stem Cell Array^®^ team**

Lauren Bauer^1^, Katie Brenner^1^, Geoff Buckley-Herd^1^, Matthew Butawan^1^, John Cerrone^1^, Anthony Chan^1^, Sean DesMarteau^1^, Patrick Fenton^1^, Peter Ferrarotto^1^, Brodie Fischbacher^1^, Camille Fulmore^1^, Jordan Goldberg^1^, Ankush Goyal^1^, Matt Green^1^, Anna K. Hahn^1^, Jenna Hall^1^, Erin Hinch^1^, Grayson Horn^1^, Christopher J. Hunter^1^, Dillion Hutson^1^, Deanna Ingrassia^1^, Selwyn Jacob^1^, Premlatha Jagadeesan^1^, Travis Kroeker^1^, Gregory Lallos^1^, Mike Leibold^1^, Hector Martinez^1^, Barry McCarthy^1^, Paul McCoy^1^, Connor McKnight^1^, Dorota N. Moroziewicz^1^, Enya O’Connor^1^, Reid Otto^1^, Temi Oyelola^1^, Katie Reggio^1^, Michael Santos^1^, Ana Sevilla^1^, Dong Woo Shin^1^, Vadim Solomonik^1^, Bruce Sun^1^, Nicole Tamarov^1^, Rebecca Tibbetts^1^, Farah Vejzagic^1^, Daniel White^1^, Christopher M. Woodard^1^, Hongyan Zhou^1^, Matthew Zimmer^1^

1. New York Stem Cell Foundation Research Institute, New York, NY 10019, USA

**GENESiPS consortium**

Ivan Carcamo-Orive^1,2^, Gabriel E. Hoffman^3^, Paige Cundiff^4^, Noam D. Beckmann^3^, Sunita L. D’Souza^5^, Joshua W. Knowles^6,7,8,9^, Achchhe Patel^4^, Dimitri Papatsenko^4,10^, Fahim Abbasi^11^, Gerald M. Reaven^11^, Sean Whalen^12^, Philip Lee^11^, Mohammad Shahbazi^11^, Marc Y. R. Henrion^3^, Kuixi Zhu^3^, Sven Wang^3^, Panos Roussos^3,13,14^, Eric E. Schadt^3^, Gaurav Pandey^3^, Rui Chang^3^, Thomas Quertermous^11^, Ihor Lemischka^5^

1. IKERBASQUE, Basque Foundation for Science, Bilbao 48009, Spain

2. Department of Endocrinology, Metabolism, Nutrition, and Kidney Disease, Biobizkaia Health Research Institute, Cruces 48903, Spain

3. Department of Genetics and Genomic Sciences, Institute of Genomics and Multiscale Biology, Icahn School of Medicine at Mount Sinai, New York, NY 10029, USA

4. Department of Developmental and Regenerative Biology, Black Family Stem Cell Institute, Icahn School of Medicine at Mount Sinai, New York, NY 10029, USA

5. Department of Developmental and Regenerative Biology, Experimental Therapeutics Institute, Black Family Stem Cell Institute, Icahn School of Medicine at Mount Sinai, New York, NY 10029, USA

6. Department of Medicine, Division of Cardiovascular Medicine, Stanford University School of Medicine, Stanford, CA 94304, USA

7. Stanford Cardiovascular Institute, Stanford University School of Medicine, Stanford, CA 94304, USA

8. Stanford Diabetes Research Center, Stanford University School of Medicine, Stanford, CA 94304, USA

9. Stanford Prevention Research Center, Stanford University School of Medicine, Stanford, CA 94304, USA

10. Skolkovo Institute of Science and Technology, Nobel Street, Building 3, Moscow 143026, Russia

11. Department of Medicine and Cardiovascular Institute, Stanford University School of Medicine, Stanford, CA 94305, USA

12. Gladstone Institutes, University of California, San Francisco, San Francisco, CA 94148, USA

13. Department of Psychiatry, Icahn School of Medicine at Mount Sinai, New York, NY 10029, USA

14. Mental Illness Research, Education, and Clinical Center (VISN 3), James J. Peters VA Medical Center, Bronx, NY 10468, USA

**iPSCORE consortium**

Lana R. Aguiar^1^, Angelo D. Arias^2^, Timothy D. Arthur^3,4^, Paola Benaglio^2^, W. Travis Berggren^5^, Juan C. I. Belmonte^6^, Victor Borja^1^, Megan Cook^1^, Matteo D’Antonio^4^, Agnieszka D’Antonio-Chronowska^7^, Christopher DeBoever^8^, Kenneth E. Diffenderfer^5^, Margaret K. R. Donovan^4,8^, KathyJean Farnam^1^, Kelly A. Frazer^1,9^, Kyohei Fujita^9^, Melvin Garcia^1^, Olivier Harismendy^4^, Benjamin A. Henson^1,9^, David Jakubosky^3,4^, Kristen Jepsen^1^, Isaac Joshua^1^, He Li^9^, Hiroko Matsui^1^, Angelina McCarron^1^, Naoki Nariai^9^, Jennifer P. Nguyen^4,8^, Daniel T. O’Connor^10^, Jonathan Okubo^1^, Athanasia D. Panopoulous^11,12^, Fengwen Rao^10^, Joaquin Reyna^1^, Bianca M. Salgado^1^, Nayara Silva^9^, Erin N. Smith^9^, Josh Sohmer^1^, Shawn Yost^8^, William W. Y. Greenwald^8^

1. Institute of Genomic Medicine, University of California San Diego, La Jolla, CA 92092, USA

2. Department of Pediatrics, University of California, La Jolla, CA 92092, USA

3. Biomedical Sciences Program, University of California, San Diego, La Jolla, CA 92092, USA

4. Department of Biomedical Informatics, University of California, San Diego, La Jolla, CA 92092, USA

5. Stem Cell Core, Salk Institute for Biological Studies, La Jolla, CA 92092, USA

6. Altos Labs, Inc., San Diego, CA 92121, USA

7. Center for Epigenomics, University of California San Diego, School of Medicine, La Jolla, CA 92092, USA

8. Bioinformatics and Systems Biology Graduate Program, University of California, San Diego, La Jolla, CA 92092, USA

9. Department of Pediatrics, University of California, San Diego, La Jolla, CA 92092, USA

10. Department of Medicine, University of California, San Diego, La Jolla, CA 92092, USA

11. Board of Governors Regenerative Medicine Institute, Cedars-Sinai Medical Center, Los Angeles, CA 90048, USA

12. Department of Biomedical Sciences, Cedars-Sinai Medical Center, Los Angeles, CA 90048, USA

## Methods

### EXPERIMENTAL MODEL AND STUDY PARTICIPANT DETAILS

#### Summary of induced pluripotent stem cells (iPSCs)

We used existing lines and newly generated lines. We obtained existing lines from four primary studies: the Human Induced Pluripotent Stem Cells Initiative (HipSci; n = 33),^22^ GENEticS of Insulin Sensitivity (GENESiPS; n = 119),^26^ iPSC Collection for Omic REsearch (iPSCORE; n = 183),^23,25^ and NYSCF Global Stem Cell Array (GSA; n = 255).^24^ In addition, we generated 12 iPSC lines from human fibroblasts of consented donors from the Finland-United States Investigation of NIDDM (FUSION) study^21^ using the NYSCF automated, high-throughput iPSC characterization and differentiation platform as described previously.^24^ In total, this study considered 602 lines, which included sixteen twin participants (eight twin pairs).

#### iPSC thawing and processing

As iPSCs were delivered to NYSCF in various non-standard, non-uniform cryovials, they were not immediately available for thawing. As such, we used manual processes to thaw iPSCs not stored in Matrix tubes. We removed cryovials containing frozen iPSCs from liquid nitrogen storage and transferred them on dry ice before thawing. We placed vials into a 37°C water bath for ∼90 seconds before resuspending them in prewarmed mTeSR1 medium (STEMCELL Technologies, Cat: #85850, Vancouver, BC) containing 10% CloneR (STEMCELL Technologies, Cat: #05888). We transferred cells into a 15 mL conical tube and added additional mTeSR1 with 10% CloneR dropwise to the conical tube to ensure complete neutralization of the cryopreservative medium for a total volume of 15 mL per conical tube. We took a small aliquot (10 µL) of each cell suspension for cell counting using a dead/total count application on an Opera Phenix High-Content Screening System (herein referred to as Opera Phenix; PerkinElmer, Waltham, MA). We centrifuged cell suspensions at 800 revolutions per minute for four minutes and aspirated the supernatant without disturbing the cell pellet. We resuspended cells in mTeSR1 with 10% CloneR media solution and plated cells on a 12-well plate, 24-well plate, or 96-well Geltrex coated plate (Thermo Fisher, Cat: #A1413301), depending on their respective live cell counts as follows. If we counted fewer than 100,000 live cells for a given vial, we resuspended cells in 200 µL of mTeSR + 10% CloneR media and seeded cells into one well of a 96-well plate; if we counted fewer than 250,000 but more than 100,000 live cells, we resuspended cells in 500 µL of media solution and seeded cells into one well of a 24-well plate; and if we counted over 250,000 live cells, we resuspended cells in 1.0 mL of media solution seeded into one well of a 12-well plate.

#### iPSC culture, expansion, and passaging

After thawing and plating, we cultured cells for 48 hours at 37°C in 5% CO_2_ and mTeSR1 with 10% CloneR without changing media and without penicillin-streptomycin. We fed cells daily with mTeSR1. We collected supernatant from each well and tested for mycoplasma contamination with the MycoAlert® Mycoplasma PLUS Detection Kit (Lonza, Cat: #LT07-710, Rockville, MD) and the accompanying MycoAlert® Assay Control Set (Lonza, Cat: #LT02-518) as per the manufacturer’s instructions. After we tested for mycoplasma contamination, we transferred mycoplasma-free cells into the automated workcells and fed them mTeSR1 supplemented with penicillin-streptomycin (pen-strep; Thermo Fisher, Cat: #15140122). Upon reaching 60-80% confluence, we passaged cells on the NYSCF robotic culture system into a Geltrex coated 24-well plate (Thermo Fisher) at a density of 50,000 cells per well. We performed all liquid handling steps using an OT-2 liquid handler (OpenTrons, Long Island City, NY) and took measurements on a Synergy HT BioTek plate reader (BioTek, Winooski, VT).

We aspirated culture medium without disturbing the cell layer and washed wells with Accutase (Sigma-Aldrich, Cat: #A6964, St. Louis, MO). We dissociated cells in this same reagent before performing additional washes with 250 µL of Accutase. We incubated cells for 20 minutes at 37°C in 5% CO_2_. Once cells were fully dissociated, we neutralized Accutase by adding prewarmed mTeSR1 containing 10 µM Y27632 (Accutase-neutralization media ratio: 1:1). We added 100 µL of mTeSR1 for a 96-well plate; 250 µL of mTeSR1 for a 24-well plate, and 500 µL of mTeSR1 for a 12-well plate. We transferred cell suspensions into an intermediate 96-well round-bottom plate and centrifuged at 120 g for five minutes. After centrifugation, we aspirated the supernatant and resuspended cells in 1.0 mL of prewarmed mTeSR1 containing 10 µM of Y27632. We took 10 µL of each cell suspension for a dead/total cell count on the Opera Phenix. We transferred cells from the intermediate plate to the destination plate(s) at the appropriate cell density (for 24-well plates, 50,000 live cells per well; for 96-well plates, 10,000-20,000 live cells per well). We then returned destination plates to the incubator for a minimum of 24 hours.

After 24 hours, we performed a half-feed followed by daily media changes. Upon reaching 60-80% confluence, we passaged cells into Geltrex coated 96-well plates (Thermo Fisher) with 10,000-20,000 cells per well in mTeSR1 with pen-strep and 10 µM Y27632 dihydrochloride (AbMole, Cat: #M1817, Houston, TX). After 24 hours, we performed a half-feed before subsequent daily feeds with mTeSR. Upon reaching ∼90% confluency, we dissociated and counted cells before we transferred the cell suspension into Matrix tubes and centrifuged at 600×g for 5 minutes. We aspirated the supernatant without disturbing the cell pellets. We resuspended cells intended for bulk RNA sequencing in 100 µL of TRIzol™ (Thermo Fisher; Cat: #15596026) per Matrix tube and stored at −80°C. We left cells intended for SNP genotyping and genome/exome sequencing as dry pellets.

#### Ethics statement

All research in this study was conducted under category 1A as defined by the International Society for Stem Cell Research guidelines. All tissue samples were sourced under IRB #: OH95-HG-N030. Study subjects provided tissues under full consent and agreement to generate and use iPSC lines for broad research. All samples are anonymized such that personally identifiable information is not accessible.

### METHOD DETAILS

#### RNA extraction and sequencing

We performed bulk RNA sequencing for all cell lines in-house at the NIH Intramural Sequencing Core (NISC). We extracted total RNA by transferring the 100 µL TRIzol™ cell suspension to a 1.5 mL microfuge tube and adding 900 µL of additional TRIzol™ following the manufacturer’s instructions. We eluted the RNA in 20 µL RNAse-free water. We assessed total RNA using the RNA 6000 Nano Kit (Agilent, Cat. #5067-1511, Santa Clara, CA) and quantitated using the QubitTM RNA Broad Range Assay kit (Thermo Fisher, Cat. #Q10211). We used 1.0 µg of total RNA with a RNA Integrity Number (RIN) ≥ 7 for library preparation with the NEBNext Poly(A) mRNA Magnetic Isolation Module (New England Biolabs, Cat. #E7490, Ipswich, MA). We sequenced libraries on the Illumina NovaSeq 6000 platform using a S4 flow cell with 151 bp paired-end reads.

#### DNA extraction, genotyping, and sequencing

AKESOgen (Tempus, Chicago, IL) performed SNP genotyping of all NYSCF GSA iPSC lines. We extracted DNA using the Promega Maxwell 96 gDNA Miniprep HT System following the manufacturer’s recommendations (Promega, Cat. #A2670, Madison, WI). We transferred DNA to AKESOgen for sequencing using the Illumina Global Screening Array and processed raw data using Illumina Genome Studio with reference genome GRCh37. We used three Illumina BeadChip arrays: InfiniumOmni2-5Exome-8v1-3 v1.3 (Illumina, Cat. #20031813, San Diego, CA), Infinium Global Screening Array-24 Kit (Illumina, Cat. #20030770), and Infinium HumanCore-24 v1.2 (Illumina, Cat. #20024566).

We performed SNP genotyping of all other iPSC lines (i.e., from the FUSION, HipSci, iPSCORE, and GENESiPS cohorts) at the NHGRI Genomics Core. We resuspended DNA pellets in 100 µL 1X PBS (Thermo Fisher, Cat. #10010031) and transferred to a 1.5 mL microfuge tube (Fisher Scientific, Cat. #14-222-169). We added 100 µL of additional 1X PBS to wash the Matrix tubes and combined the volumes. We extracted DNA using the QiaAMP Mini kit (Qiagen, Cat. #51304, Hilden, Germany) following manufacturer instructions. We resuspended DNA in 50 µL Buffer AE and quantitated using QubitTM 1X dsDNA Broad Range Assay Kit (Thermo Fisher, Cat. #Q33266). We genotyped 250 ng of DNA using the Ilumina Omni 2.5 Exome-8 v1.5 beadchip array. We analyzed data in GenomeStudio with the v1.5_A1_GRCh37 manifest and the v1.5_A1 cluster file (Illumina).

Genosity (Invitae Corporation, South San Francisco, CA) performed exome sequencing (ES) of NYSCF GSA iPSC lines. We extracted DNA using the Promega Maxwell 96 gDNA Miniprep HT System following manufacturer’s instructions (Promega). We transferred DNA to Genosity for sequencing using Twist Human Core Exome EF Multiplex Complete Kit (Twist Bioscience, Cat. #PN 100803, South San Francisco, CA). Genosity sequenced at a minimum coverage of 25X of 90% of bases for 183 samples and returned the trimmed FASTQ files for downstream analyses.

We performed genome sequencing (GS) of iPSC lines from the FUSION, HipSci, and GENESiPS cohorts at NISC. We resuspended DNA pellets in 100 µL 1X PBS (Thermo Fisher) and transferred the contents to a 1.5 mL microfuge tube (Fisher Scientific). We added 100 µL of additional 1X PBS to wash the matrix tubes and combined the volumes. We extracted DNA using the QiaAMP Mini kit (Qiagen). We resuspended DNA in 50 µL Buffer AE and quantitated using the Qubit^TM^ 1X dsDNA Broad Range Assay Kit (Thermo Fisher, Cat. #Q33266). We submitted 1.0 µg of DNA to NISC for library preparation with the Illumina TruSeq PCR-free Sample Preparation kit (Illumina, Cat. #20015962). We sequenced libraries on the Illumina NovaSeqX platform using 25B flow cells.

We downloaded genome sequencing data sourced from donor blood or fibroblasts from iPSCORE (dbGaP accession number phs001325.v6.p1).^23,25^

#### Cellular imaging of individual wells

To derive a growth rate phenotype, we used daily measurements of well coverage (expressed as a percent) for each well. We imaged all plates stored within the automated system nightly using Celigo imagers during culture (Revvity Inc., Cat. #200-BFFL-5C, Waltham, MA). We used the default confluence analysis parameters from the Celigo software to measure well coverage.

### QUANTIFICATION AND STATISTICAL ANALYSIS

#### Growth rate derivation from cellular images and replicate concordance

We only considered wells that contained at least three well coverage measurements between 30-70%. We removed outlier wells using magnitude-shape plot outlier detection in scikit-FDA v0.10.1.^49^ Because most donors had multiple replicate wells, we identified and removed wells with low concordance by calculating the pairwise Pearson correlation coefficients among all replicate wells for each donor. Specifically, we rounded all well coverage measurement time points to the nearest day and aligned the resulting time series of well coverage measurements for each well pair based on shared time points. We calculated Pearson correlation coefficients using all complete, aligned observations. We excluded any well that exhibited a Pearson correlation coefficient (r) < 0.75 with any other replicate well from the same donor. From these filtered data, we min-max normalized the individual per-well time series measurements and fitted each to a standard logistic curve (time measured in days versus normalized well coverage) using Growthcurver v0.3.1.^50^ We further excluded replicate wells that either failed to fit a growth curve or showed a residual sum of squares from the logistic curve fit exceeding three standard deviations above the mean error. Finally, for each remaining well, we derived the growth Area-Under-the-Curve (gAUC) metric from the fitted curves.

To assess concordance of gAUC across replicate wells corresponding to each donor and within the eight twin pairs, we calculated the intraclass correlation coefficient (ICC) using a one-way random-effects model for single measurements (ICC(1,1)) as implemented in the Python statsmodels package v0.14.4.^51^

#### SNP genotype quality control procedures and imputation

After SNP chip genotyping, we aligned array probe sequences to the GRCh37 reference genome using the Burrows-Wheeler Aligner (BWA) v0.17.7.^52^ Pre-imputation quality control procedures included removing multi-mapping probes, probes containing known 3’ variants, and non-SNP variants. We further filtered out variants with per-variant missingness > 5% and Hardy-Weinberg equilibrium (HWE) *P* < 10⁻⁶, and removed samples exceeding 5% missingness. Using the 1000 Genomes Project Phase 3 data as a reference,^27^ we verified and corrected strand orientation, allele assignments, genomic positions, and reference/alternate designations. We excluded biallelic variants with minor allele frequencies (MAF) > 0.4, variants absent from the reference panel, and variants with discordant allele assignments. We merged genotyping data from individual arrays prior to lifting over and imputation.

We conducted lift over from GRCh37 to GRCh38, pre-imputation variant-level quality control procedures, and genotype imputation using the TOPMed Imputation Server.^53^ We used Eagle v2.4^54^ to perform SNV pre-phasing and minimac4^53^ to impute SNV genotype dosages with the TOPMed imputation panel v3. After imputation, we retained biallelic SNPs meeting the following criteria: MAF > 0.05, imputation quality r^2^ > 0.3, and HWE *P* > 10⁻^10^. We merged the quality-controlled imputed data across studies and used PLINK v2.00a5^55^ with the “--king-cutoff” flag to remove genetically identical samples (KING relatedness coefficient > 0.354).^56^ Our final study set contained 602 unique individuals. To infer donor sex for all 602 individuals based on genotype calls, we used bcftools v1.17^57^ with the “guess-ploidy” plugin considering genotypes in the non-pseudoautosomal region of chromosome X from quality-controlled common variant data.

We also generated linkage disequilibrium (LD)-pruned genotypes for downstream analyses, including genetic principal component (PC) calculation and power analyses. We retained only biallelic SNPs with MAF > 0.05, HWE *P* > 10^−10^, and imputation quality r^2^ > 0.8. We excluded high-LD regions defined by Anderson et al.^58^ We applied LD pruning using a sliding window of 1 kb, a step size of 0.8, and an LD threshold of r^2^ < 0.1, retaining 66,917 pruned variants for downstream analysis.

#### Variant calling and quality control procedures in genomic sequencing

After GS and ES, we processed reads according to the GATK Best Practices pipeline for variant discovery.^59^ Following adapter trimming, we aligned paired-end reads to the GRCh38 reference genome using BWA MEM v0.17.7^52^ with default parameters. We subsequently filtered per-lane BAM files to retain high-quality, primary alignments, merged, and sorted using samtools v1.22.^60^ To ensure accurate variant calling, we recalibrated base quality scores using the Genome Analysis Toolkit (GATK) v4.0.5.1 BaseRecalibrator,^61^ incorporating dbSNP v151^62^ as well as the Mills and 1000 Genomes gold standard indels^59^ to correct systematic biases in base quality scores. We performed variant calling per sample in GVCF mode using the GATK HaplotypeCaller.^61^ We calibrated variants (indels and SNPs) via Variant Quality Score Recalibration (VQSR),^61^ which was trained using the Mills and 1000 Genomes gold standard indels,^59^ HapMap v3.3,^63^ sites found to be polymorphic on the Illumina Omni 2.5M SNP array,^59^ and dbSNP v151^62^ to finalize high-quality, recalibrated SNP and indel calls.^59,62,63^

We used VerifyBamID v1.1.1^64^ to compare the sequencing data to quality-controlled genotypes and identified no sample swaps. For downstream analyses, we retained biallelic variants (SNPs and indels with length ≤ 5 bp) meeting the following criteria: MAF < 0.1 and HWE *P* > 10^−10^. We kept genetically distinct samples (KING relatedness coefficient < 0.354) as determined from SNP genotype data (n = 514).

#### RNA-seq data processing and quality control procedures

After RNA sequencing, we performed transcript quantification using Salmon v1.10.3^65^ in quasi-mapping mode with default parameters, aligning reads to the GRCh38 human reference transcriptome (Ensembl v111^66^). We used tximport v1.32.0^67^ to convert transcript-level abundance estimates to gene-level counts for differential gene expression analysis via the lengthScaledTPM function.

As an additional quality control procedure, we used STAR v2.7.11a^68^ to align reads to the GRCh38 reference genome in a paired-end fashion. We identified sample swaps and contaminated samples by comparing the allelic RNA-seq read count distribution to the finalized genotypes of the 602 donors using VerifyBamID v1.1.1.^64^ We identified and resolved 38 pairs of sample swaps. We removed 25 contaminated samples from the analyses (FREEMIX or CHIPMIX > 0.055). After these procedures, 333 iPSC lines remained for final analyses. For downstream analyses, we considered only genes with transcripts per million (TPM) > 0.5 in at least 50% of samples, resulting in 15,603 features.

#### Determination of covariates for downstream analyses

To determine covariates for subsequent models, we fit a linear mixed model implemented in lmerTest v3.1.358^69^ to test if genetically-imputed donor sex, pluripotency reprogramming method, starting cell type, and cell culture media are associated (FDR < 5%, Benjamini-Hochberg procedure^70^) with gAUC, controlling for donor as a random effect. We found pluripotency reprogramming method, starting cell type, and cell culture media were associated with gAUC and therefore included these three terms in subsequent association analyses. We did not include cell culture media type as a covariate in our differential expression analyses because all cell lines included in those analyses were cultured in mTeSR media.

#### Differential expression analyses

We normalized gAUC using inverse rank normalization and calculated the gAUC mean value for each RNA-seq sample. We performed differential expression analysis using a three-stage process as previously described in Taylor et al.^71^ First, we tested for differentially expressed genes across gAUC using DESeq2 v1.36.0^72^ with default parameters and adjusting for starting cell type and pluripotency reprogramming method as fixed effect covariates. Next, we used RUVSeq v1.30.0^73^ with default parameters to derive a factor representing unwanted, latent variation using control gene expression, defined as genes with *P* > 0.25 from the initial differential expression analysis. Finally, we re-ran differential expression analysis and incorporated the calculated RUVSeq factor as an additional fixed-effect covariate. We performed multiple hypothesis correction using the Benjamini-Hochberg procedure.^74^

#### Genetic relatedness matrix calculation

We used GCTA v1.94.1^75^ to compute a genetic relatedness matrix (GRM) on the basis of common variant genotype data (MAF > 0.1). We used the GRM in the variance decomposition, common variant, and rare variant analyses.

#### Variance decomposition analyses

We performed variance decomposition analyses using GCTA v1.94.1^75^ at the cell line (i.e., donor) level with the mean gAUC value across all replicates per donor. We used three subsets of the available data: (i) all unique cell lines (n = 602) including cell culture media, starting cell type, and pluripotency reprogramming method as discrete covariates, (ii) all unique cell lines with bulk RNA-seq data incorporating starting cell type and pluripotency reprogramming method as discrete covariates (n = 333), and (iii) all unique cell lines with bulk RNA-seq data incorporating pluripotency markers as quantitative covariates and starting cell type and pluripotency reprogramming method as discrete covariates (n = 333). We modelled pluripotency with gene expression of four established pluripotency markers measured in TPM: *POU5F1* (ENSG00000204531), *SOX2* (ENSG00000181449), *NANOG* (ENSG00000111704), and *PODXL* (ENSG00000128567).^5,76^ For subsets (ii) and (iii), we did not include media as a discrete covariate since all cell lines with available RNA-seq data were cultured with mTeSR media.

We fit the following model:

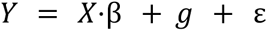

where Y represents the vector of sample-level mean phenotype values, X is the matrix of fixed-effect covariates (either media or gene expression of pluripotency markers), 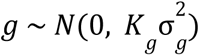 is the random genetic effect modeled using the GRM K_g_ calculated as described above, and 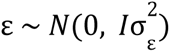 represents the residual error term. Here, 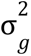 and 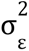 denote the genetic and residual variance components, respectively.

To quantify the proportion of phenotypic variance explained by genetic factors, we calculated the narrow-sense SNP heritability as:

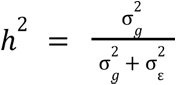

#### Genetic principal component and ancestry assignment

We calculated genetic PCs and assigned participants to genetic ancestry groups using Fast and Robust Ancestry Prediction by Online singular value decomposition and Shrinkage Adjustment (FRAPOSA)^77^. Briefly, FRAPOSA works by performing PC analysis in a diverse reference cohort with known genetic ancestry labels, systematically calculating PCs for each participant in a target cohort by comparing them to the reference, and grouping participants from the target cohort to the most similar genetic ancestry group from the reference. Using FRAPOSA with default parameters, LD-pruned genotypes as input, and 1000 Genomes Project Phase 3 data (1KG)^27^ as the reference cohort, we calculated the first 30 PCs and grouped participants into 1KG-AFR-like (African), 1KG-AMR-like (American), 1KG-EUR-like (European), 1KG-EAS-like (East Asian), or 1KG-SAS-like (South Asian) genetic ancestries.

#### Common variant association testing

We normalized gAUC growth phenotypes through rank inverse normalization. We conducted association tests using the Generalized Linear Mixed Model Association Tests (GMMAT) v1.4.2 framework with default parameters.^78^ The analyses proceeded in two steps: (i) fitting a null model using the GRM, and (ii) performing score tests to identify genetic associations. We included 10 genetic PCs, cell culture media type, starting cell type, and pluripotency reprogramming method as fixed-effect covariates. We considered variants with *P* < 5×10^−8^ as genome-wide associated and those with *P* < 10^−6^ as nominally associated.

#### Power analyses for common variant associations

We estimated the sample size needed to detect common variant associations with gAUC using an unbalanced one-way ANOVA framework, implemented in the powerEQTL package v0.3.6.^79^ Briefly, for a range of absolute effect size thresholds (absolute effect size < 0.1, 0.2, 0.3, 0.4, 0.5), we calculated statistical power for up to 6,000 samples. Calculations assumed a genome-wide threshold of 5×10^−8^, a maximum MAF of 0.5, and a residual standard deviation (σ = 0.4584) derived from variance decomposition across all 602 samples from the common variant association study. We estimated the effect size parameter “deltaVec” by averaging the observed differences in mean gAUC comparing heterozygotes to each homozygous genotype group for LD-pruned variants within each effect size bin.

#### Rare variant annotation, aggregation, and association testing

We annotated variants using Variant Effect Predictor (VEP,^80^ Ensembl v111^66^) selecting the most severe consequence across all protein-coding transcripts (Ensembl v111 annotations^66^) for each variant. We classified predicted loss of function (pLoF) variants as those with consequences including transcript ablation, splice acceptor variant, splice donor variant, stop gained, frameshift variant, stop lost, start lost, transcript amplification, feature elongation, or feature truncation. We aslo evaluated VEP-annotated missense variants for deleteriousness using CADD v1.6^81^ and classified variants with scores ≥ 20 as damaging.

For each gene, we aggregated coding variants with MAF < 0.01 that were annotated as either pLoF or damaging missense. To be considered in downstream analyses, we required a gene to contain at least two qualifying variants and carry a non-zero burden score in at least 1% of the cohort, resulting in 8,480 genes.

We normalized gAUC growth phenotypes through rank inverse normalization. We tested for associations using GMMAT v1.4.2^82^ with default parameters. The analyses proceeded in two steps: (i) fitting a null model using the GRM, (ii) performing hybrid set-based tests to identify gene-level rare variant associations using the provided variant-gene annotations. For the hybrid test, we used the SMMAT function implemented in GMMAT, which efficiently combines a burden test and an adjusted SKAT test. As covariates, we included 10 genetic PCs, cell culture media type, starting cell type, and pluripotency reprogramming method as fixed-effect covariates. We applied a Bonferroni-corrected threshold to account for the number of unique genes tested (*P* < 5.90×10^−6^, 8,480 tests). For the sensitivity analysis, we fit a regression in the same manner but using CADD ≥ 25 to define damaging missense variants, resulting in an adjusted Bonferroni-corrected threshold of *P* < 1.45×10^−5^ (3,441 tests) after gene aggregation.

## Supplemental Tables

**ESM Table S1.**
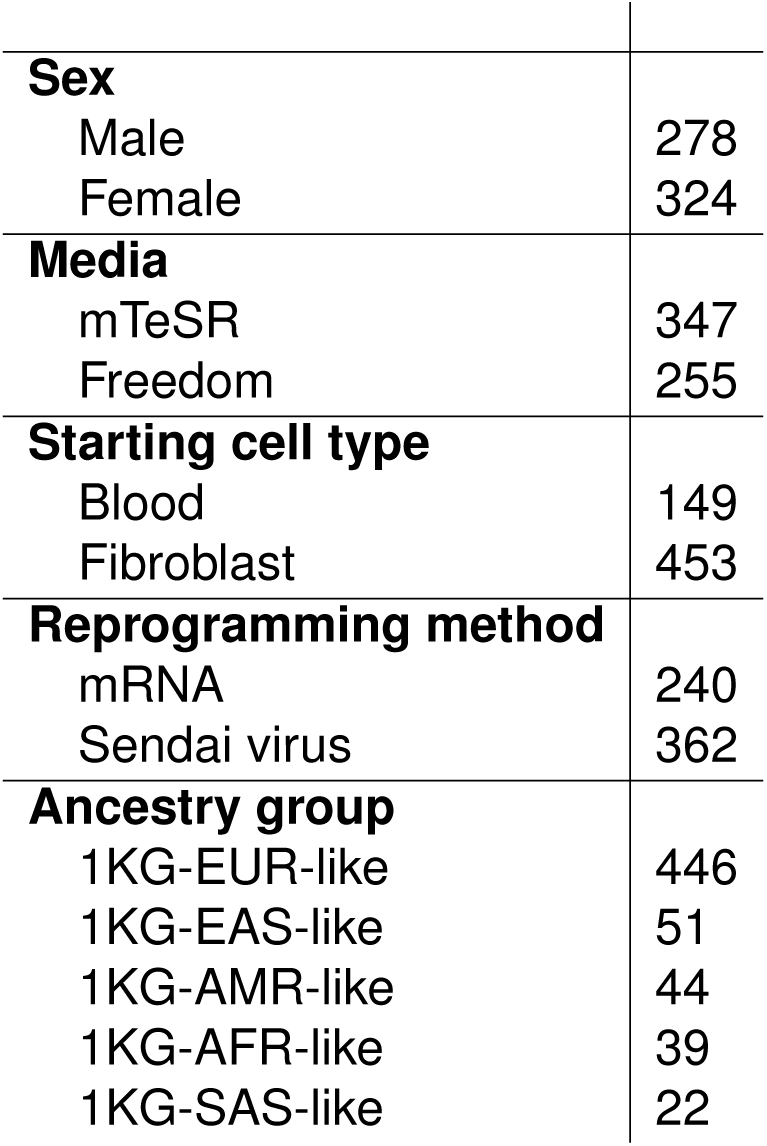
Summary statistics of 602 iPSC line attributes.

**ESM Table S2.** *WDR54* variants from the rare variant association study. Genomic coordinates from GRCh38. Source of genotype DNA reported in columns: “Blood DNA”, “Fibroblast DNA”, and “iPSC DNA.”

| rsID | Variant Location | Variant Type | CADD | Blood DNA | Fibroblast DNA | iPSC DNA |
| --- | --- | --- | --- | --- | --- | --- |
| rs147527356 | chr2:74422334 | Missense | 22.5 | 0 | 0 | 2 |
| rs150243972 | chr2:74424959 | Missense | 21.8 | 0 | 2 | 0 |
| rs187456631 | chr2:74425080 | Missense | 25.2 | 0 | 4 | 0 |
| rs139454931 | chr2:74425113 | Missense | 22.2 | 0 | 0 | 4 |
| rs140810540 | chr2:74425208 | Missense | 25.9 | 0 | 0 | 5 |

## Supplemental Figures

**ESM Fig. S1.**
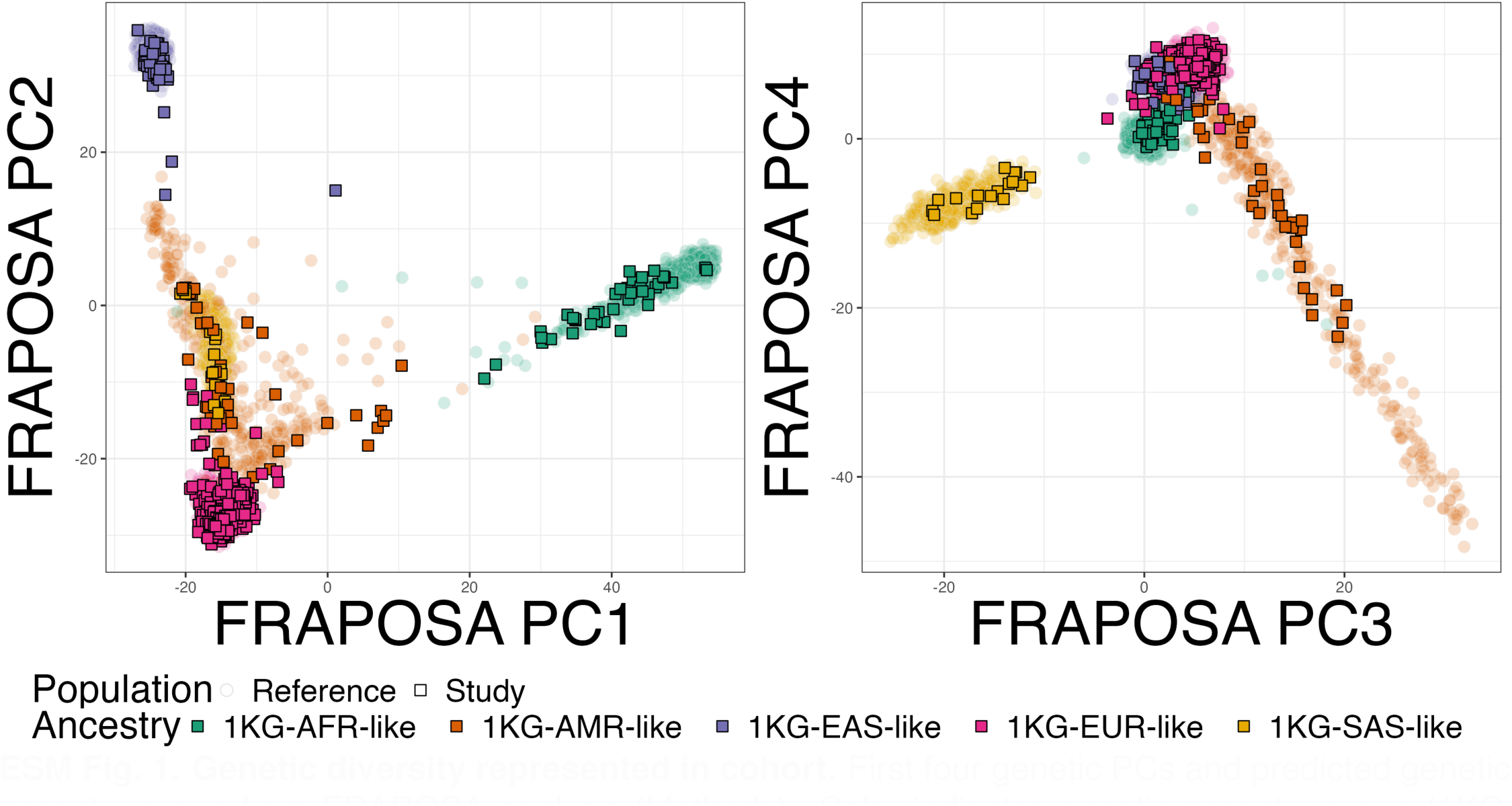
Genetic diversity represented in cohort. First four genetic PCs and predicted genetic ancestry group from FRAPOSA analysis (Methods). Color indicates genetic ancestry group (1KG-AFR-like, 1KG-AMR-like, 1KG-EAS-like, 1KG-EUR-like, or 1KG-SAS-like). Left-hand side plot displays principal components 1 versus 2; right-hand side plot displays principal components 3 versus 4. Point shape indicates study samples (squares) and reference panel samples (circles).

